# Trait genetic architecture and population structure determine model selection for genomic prediction in natural *Arabidopsis thaliana* populations

**DOI:** 10.1101/2024.07.09.601435

**Authors:** Patrick M. Gibbs, Jefferson F. Paril, Alexandre Fournier-level

## Abstract

Genomic prediction applies to a wide range of agronomically relevant traits, with distinct ontologies and genetic architectures. Selecting the most appropriate model for the distribution of genetic effects and their associated allele frequencies in the training population is crucial. Linear regression models are often preferred for genomic prediction. However, linear models may not suit all genetic architectures and training populations. Machine Learning approaches have been proposed to improve genomic prediction owing to their capacity to capture complex biology including epistasis. However, the applicability of different genomic prediction models, including non-linear/non-parametric approaches, have not been rigorously assessed across a wide variety of plant traits in natural outbreeding populations. This study evaluates genomic prediction sensitivity to trait ontology and the impact of population structure on model selection and prediction accuracy. Examining 36 quantitative traits measured for 1000+ natural genotypes of the model plant *Arabidopsis thaliana*, we assessed the performance of penalised regression, random forest, and multilayer perceptron at producing genomic predictions. Regression models were generally the most accurate, except for biochemical traits where random forest performed best. We link this result to the genetic architecture of each trait – notably that biochemical traits have simpler genetic architecture than macroscopic traits. Moreover, complex macroscopic traits, particularly those related to flowering and yield, were strongly correlated to population structure, while molecular traits were better predicted by fewer, independent markers. This study highlights the relevance of machine learning approaches for simple molecular traits and underscores the need to consider ancestral population history when designing training samples.

**Article summary:** Machine learning and linear models were tested for genomic prediction of multiple traits in the model plant *Arabidopsis thaliana*. We associate the performance of genomic prediction models to trait ontology, finding machine learning approaches applicable to biochemical traits, and linear models best for macroscopic traits. We link this result to the genetic architecture of each trait and patterns of selection in the association panel’s ancestral population, thus underscoring the relevance of these two sensitivities to genomic prediction in plant breeding.

## 1. Introduction

In plant breeding, genomic prediction can apply to any agronomically relevant traits form yield, to phenology, metabolite concentrations or the response to stresses, which all have different genetic basis. Heritable traits, whether simple and controlled by a few loci, or complex and controlled by many variants, can be predicted based on the information from dense, genome-wide markers. Genomic prediction models are often fitted to training populations designed to maximise diversity, including genotypes with varying levels of co-ancestry and different evolutionary histories (Fodor et al., 2014). Therefore, critical in genomic prediction is the choice of a model that most adequately factors population’s allele frequencies and the distribution of genetic effects for each particular trait.

The number of genome-wide markers (*p*) being typically much larger than the number of individual observations (*n*), overfitting is a major challenge in genomic prediction. A common solution is to use penalised linear regression models (Meuwissen et al., 2001). Penalised regression reduces overfitting by shrinking the effect of each marker. Most penalisation functions assume a prior distribution of the marker effects, which relates to the trait’s assumed genetic architecture. Linear models also assume that genetic factors influence the phenotype additively. This simplification allows many predictors to be incorporated parsimoniously but excludes interactive effects such as dominance and epistasis. Indeed, an additive model with a fixed prior distribution may not be optimal when the development and regulation of the focal trait are a complex function of genetic and environmental factors (Meuwissen, 2009; Morgante et al., 2018; Putra et al., 2023).

Machine learning has been proposed as a means to better accommodate biological complexity in genomic prediction (Carlborg & Haley, 2004). Non-linear, non-parametric models can account for dominance and epistasis and have fewer assumptions about the distribution of genetic effects. It is thus hypothesised that machine learning should be effective for genomic prediction under increased model complexity, as is the case for other biological problems like predicting protein structure (Jumper et al., 2021). However, the performance of non-linear machine learning models has been largely underwhelming in genomic prediction. Multiple studies have shown machine learning techniques to be no more accurate than linear regression for complex polygenic traits (Abdollahi-Arpanahi et al., 2020; Azodi et al., 2020; Bellot et al., 2018). One explanation for the under-performance of machine learning models is that the number of genetic variants used as predictors is too high compared to the number of genotypes observed in the training set. There might be many genetic interactions contributing to the development of a complex trait such as crop yield; if the effect of each interaction is small, it is difficult to parameterise them with low statistical power. Simulation studies have shown that machine learning models might rather be suited to simpler traits controlled by a small number (<100) of causal loci (Abdollahi-Arpanahi et al., 2020). Indeed, non-linear models have been most successful for predicting the occurrence of human autoimmune diseases, where most of the genetic variance can be attributed to a discrete set of loci from the major histocompatibility complex (Ohta et al., 2024). As such, rather than attempting to apply machine learning to genomic prediction generally, machine learning models might be specifically suited to simpler traits controlled by a few loci with strong epistatic effects (Abdollahi-Arpanahi et al., 2020; Farooq et al., 2023). This hypothesis is agronomically relevant. In crops, it can be desirable to breed for biochemical traits, for example the concentration of minerals or metabolites (Šimić et al., 2012), which often have simple genetic architectures (Riedelsheimer et al., 2012). On the other hand, macroscopic traits such as those related to yield often combine many endophenotypes and tend to be more complex.

In addition to biological complexity, different evolutionary histories and patterns of selection in the ancestral populations of training panels could also affect model selection (Fodor et al., 2014; Guo et al., 2014). Population-specific selection on a trait leads to covariance between trait value and population structure, confounding variants causally affecting a trait and background genomic variations captured by neutral markers. Thus within one training panel, traits may be confounded to a different degrees by historical patterns of selection. Moreover, the level of selection on a trait, the trait’s phenotypic ontology, and its architecture are likely to be linked, making it extremely arduous to obtain independent estimates of each factor (Zhang, 2018). For example, flowering time in *A. thaliana* is known to be both complex in genetic architecture and selected along environmental clines (Zan & Carlborg, 2019; Fournier-Level et al., 2022).

The design and tuning of genomic prediction studies requires an understanding of how genetic architectures and evolutionary histories interact to produce trait variation in the training population considered. However, there has been very limited sensitivity analyses comparing linear and machine learning models across a wide range of genetic architectures in natural, outbreeding plant populations. Our study aims to address this gap by comparing the accuracy of machine learning and linear models for the prediction of 36 *Arabidopsis thaliana* traits. We specifically test the sensitivity to the phenotypic ontology of each trait, the genetic architecture of each trait and finally assess how population structure and demography affect predictions. We investigate how intertwined these three factors are, testing the hypothesis that linear models would perform best for macroscopic traits, with complex genetic architecture that is structured along environmental gradients. On the other hand, machine learning approaches are expected to improve prediction accuracy for biochemical traits, which usually have a simple genetic architecture, and are less heterogeneously selected across environments.

## 2. Methods

### 2.1 Data Set

We used 10,709,466 molecular markers genotyped across 2029 inbred homozygous *Arabidopsis thaliana* accessions from the ‘K2029’ public data set (Arouisse et al., 2020) which combines genotypes from the RegMap Panel (Horton et al., 2012) and the 1001 genome project (Alonso-Blanco et al., 2016). We downloaded all available phenotypes measured in all or a subset of these accessions from the AraPheno database (Seren et al., 2017). In addition to the traits listed in the AraPheno database, we also included 22 metabolic traits, which are the concentration of glucosinolates involved in *Arabidopsis thaliana*‘s herbivory stress response (Brachi et al., 2015). No phenotype was measured across all genotyped accessions, meaning experiments always had less than 2029 observations. For traits where measures were replicated within an accession, we used the breeding value estimated across the replicates as the trait value for our genomic prediction experiments. We estimated these breeding values using the best linear unbiased predictor (BLUP). Using a phenotypic value derived from replicated measures of the same accession minimises the effect of the environment in the study. From all traits available in the AraPheno database, we retained only studies with 150 or more phenotyped accessions. When two traits were correlated with Pearson’s correlation *r*^2^ > 0.75, we arbitrarily retained one, and subsequently only considered traits where at least one genomic prediction model could explain 20% of the phenotypic variance. This filtering process reduced the set of traits from 551 to 36. We then grouped these 36 traits by their associated trait ontology terms (Cooper et al., 2024).

The total number of molecular markers was filtered to keep computation time within a practical range. For each trait, markers with minor allele frequencies (MAF) lesser than 5% were discarded. We used ‘plink thin’ to prune correlated markers within a sliding window along the genome, using a window size of 200 kbp, removing one of each pair of markers with linkage disequilibrium *r*^2^ > 0.6. Due to each trait having a different set of phenotyped accessions, we conducted this filtering strategy independently for each trait. The total number of markers used for the final set of 36 traits ranged from 218,128 to 366,534.

### 2.2 Analysis of Genetic Architecture

We investigated the genetic architecture of each trait using genome-wide association studies (GWAS). GWAS were conducted using the ‘VCFtoGWAS’ function (Vogt et al., 2022) implementing a ‘GEMMA’ model (Zhou & Stephens, 2012), where the effect of pairwise relationships (estimated as identity-by-state-based kinships) is modelled as random, and the effect of individual markers are modelled as fixed and tested using a Wald test. Polygenic traits were defined when showing a high heritability while lacking substantial GWAS peaks; oligogenic traits where defined as displaying multiple GWAS peaks; and monogenic traits were defined when displaying a single associated locus (Atwell et al., 2010). The trait complexity classification defined above was determined quantitatively by the kurtosis of the distribution of GWAS p-values, and qualitatively validated through the inspection of Manhattan plots.

### 2.3 Prediction Models

#### 2.3.1 Penalised Linear Regression

For the linear models, we implemented Ridge regression/GBLUP (genomic best linear unbiased predictor), Lasso, and Elastic-net. Briefly, these models fit a weighted linear combination of regression coefficients for the predictors (denoted ***β***) by minimising the following objective function:

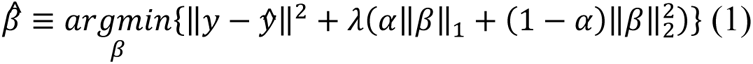

Where *y* is a vector of observed phenotypes and *ŷ* is the predicted phenotype, γ controls the penalisation and α weights L_1_ and L_2_ penalisation. When α is null, the model is equivalent to Ridge regression, when α is 1 the model is equivalent to Lasso regression, and when 0 < α < 1 the model is termed Elastic-net regression (Tibshirani, 1996; Zou & Hastie, 2005).

In our implementation of these models, the meta-parameters γ, and α in the case of Elastic-net, were chosen through five-fold cross-validation in the training partition of the dataset searching the following domains: γ ∈ {1.5^−30^…1. 5^30^}, α ∈ {0.01,0.1,0.5,0.85,0.9,0.97,0.99,0.995}. Ridge and Lasso regressions were implemented using the ‘RidgeCV’ and ‘LassoCV’ functions of the scikit-learn package (Pedregosa et al., 2011). Elastic-net regression was implemented using ‘glmnet’ with the coordinate descent algorithm (Friedman et al., 2010). All other parameters were kept to default values.

#### 2.3.2 Multilayer Perceptron

A Multilayer Perceptron (MLP) is a simple ANN model where perceptrons are organised in fully connected layers. In our implementation, we used a rectified linear unit (ReLU) activation function. Preliminary tests showed that other sigmoidal activation functions (such as *tanh*) had similar prediction accuracy. We optimised {1,2,3} hidden layers in the network, where the number of perceptrons in each hidden layer was set to two-thirds of the number of input predictors. We also incorporated L2 penalisation where γ is a scalar optimised over {0, 0.01, 0.1, 1}. Weights were fitted iteratively using the Adam optimiser (Kingma & Ba, 2014). To determine when to cease the training of the neural network model, we used an early stopping approach (Prechelt, 1998). Briefly, we held out 10% of observations, and training was stopped when either out of sample mean squared error did not decrease by at least the tolerance value for 10 consecutive epochs (iterations over the dataset), or 250 epochs were reached. The tolerance value was optimised over {0.05, 0.01, 0.005}, noting here that each phenotype was standardised to have a variance of 1. We also added dropouts (Srivastava et al., 2014). Dropouts are a type of regularisation specifically designed for ANNs which randomly deactivate, or drop out, a certain percentage of weights between layers during each training iteration. This prevents the model from becoming too reliant on any specific perceptron or connection. We optimised the network either using no dropout or dropping out 10% of the network connections during training. Due to the computationally heavy requirement of fitting ANNs, we applied an additional filtering to the number of markers used to fit this model. For this filtering, we fitted a linear Lasso regression then filtered out markers with a regression coefficient equal to 0. Here, the penalty terms were determined via five-fold cross-validation alongside the other meta-parameters specified in this section.

#### 2.3.3 Random forest

Random forest is an ensemble model that fits decision trees to bootstrapped samples of the data, then makes predictions by averaging the output of each tree. We implemented the ‘RandomForest’ sci-kit learn function (Pedregosa et al., 2011). Five-hundred decision trees were used across experiments; increasing the number of decision trees did not affect performance. Each of these trees was fitted to the full set of predictors, randomness between trees being introduced through bootstrapping of the accessions. All other parameters were kept as default.

### 2.3 Model benchmarking and meta-parameter selection

We compared the accuracies and residual errors of each of the models described using nested 10-fold cross validation. This framework consists of an inner loop and an outer loop. The outer loop is a canonical ten-fold cross-validation and measures generalizability. The inner loop takes each training partition from the outer loop and uses five-fold cross validation to evaluate each meta-parameter configuration of the tested model.

### 2.4 Analysis of Spatial Clines

To analyse the effect of differential selection across genotypes having evolved across distant environments, we quantified the strength of spatial clines for each trait. To do this, we used the following model of the phenotypic values in terms of the latitude and longitude of where each accession was originally sampled from.

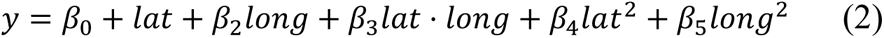

We quantified the strength of a spatial cline based on the phenotypic variance that was captured by this model. We applied this model only to accessions sampled within Europe (latitude -40–60, longitude 20–70), excluding the small number of accessions sampled from North America.

### 2.5 Code Availability

All code and the corresponding analysis used in this paper can be found on the following repository: https://github.com/Patrick-Gibbs/Trait-genetic-architecture-and-population-structure-genomic-prediction

## 3. Results

### 3.1 Optimal genomic prediction model is associated with trait ontology

Of the 551 publicly available traits, 36 traits were independent (correlation between traits *r*^2^< 0.75) and sufficiently heritable to be predicted with an accuracy *r*^2^ > 0.2 by one of the genomic prediction models. Of this filtered set of traits, 23 were macroscopic and 13 were biochemical (Supplementary Table S1). Penalised regression yielded the most accurate predictions for all 23 macroscopic traits; 20 through Ridge regression and three through Lasso. In contrast, no single model was superior at predicting the 13 biochemical traits (Supplementary Table S2). Noticeably, Random Forest improved prediction above linear models for 7 biochemical traits (Figure 1a). The increase in accuracy was particularly large for allyl concentration, a glucosinolate involved in preventing herbivory, and molybdenum concentration, an essential micronutrient (Figure 1b, paired t-test *p* = 2.87 × 10^−4^, *p* = 3.28 × 10^−5^respectively).

**Figure 1:**
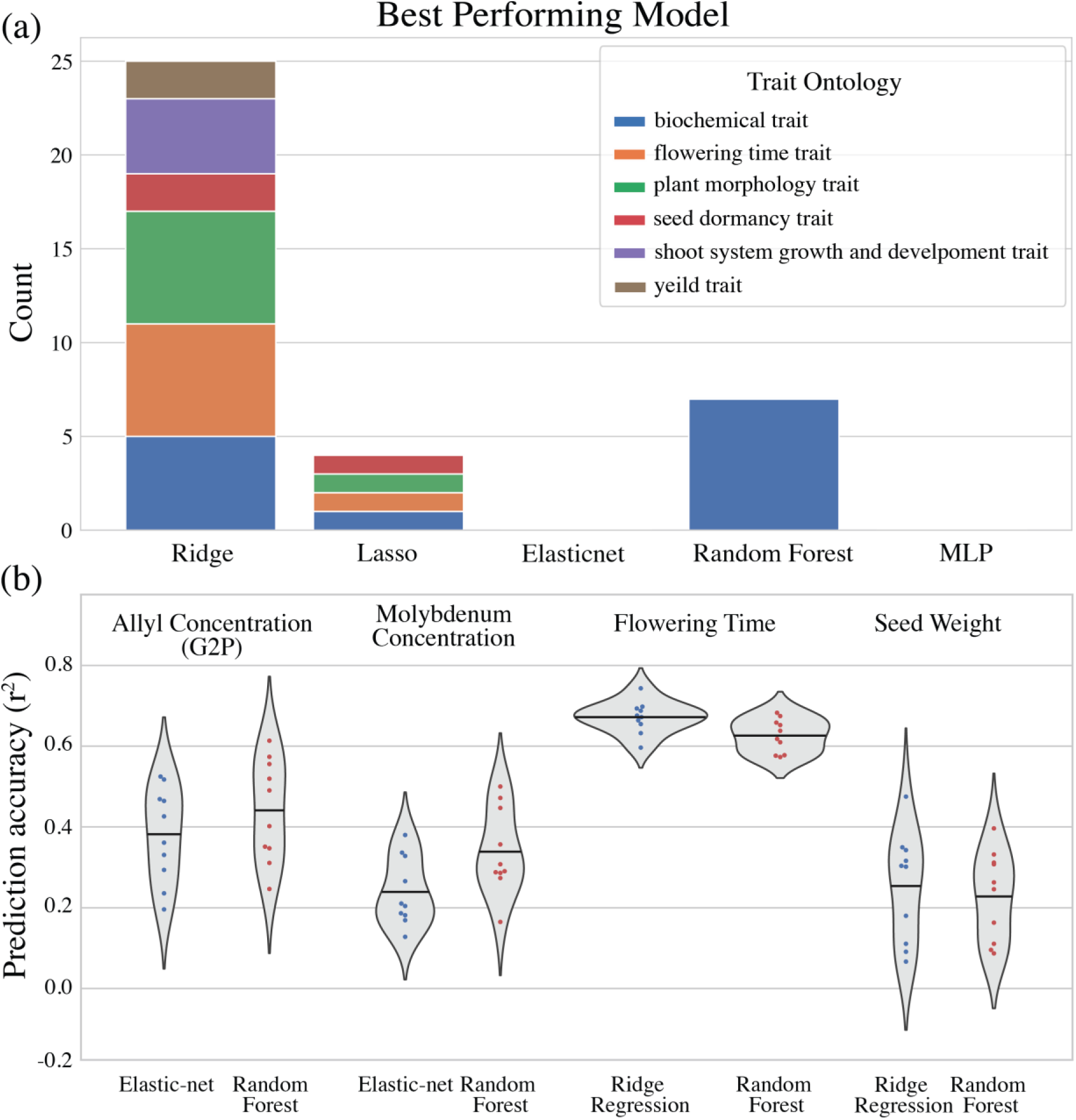
Genomic prediction accuracy depends on trait ontology class. (a) Most accurate genomic prediction model for each trait ontology term. (b) Prediction accuracy of the best linear model compared to Random Forest contrasting for a pair of biochemical traits (the concentration of allyl and molybdenum) and macroscopic traits (number of days to flowering at 10°C and average seed weight). Here, the ten data points are the prediction accuracy (measured as *r*^2^) for each partition of the ten-fold cross-validation.

### 3.2 The genetic architecture is associated with genomic prediction model choice

To test the hypothesis that random forest out-performance over penalised regression is due to the specific genetic architecture of biochemical traits, we analysed each trait through GWAS. Biochemical traits had stronger individual associations compared to macroscopic traits (Fig. 2A; Suppl. Fig. S1). Macroscopic traits generally displayed fewer association peaks despite high heritability, indicative of polygenicity. This observation was reflected in the kurtosis of the distribution of p-values, which was on average higher for biochemical traits compared to macroscopic traits (t-test, t=5.54, p=3.34e-06; Fig. 2a). Moreover, among the biochemical traits best predicted by Random Forest, six out of seven were most accurately predicted by linear models that assume sparsity (Elastic-net or Lasso). Only one of these seven traits was most accurately predicted by Ridge regression, which assumes polygenicity (Supplementary Table S2).

**Figure 2:**
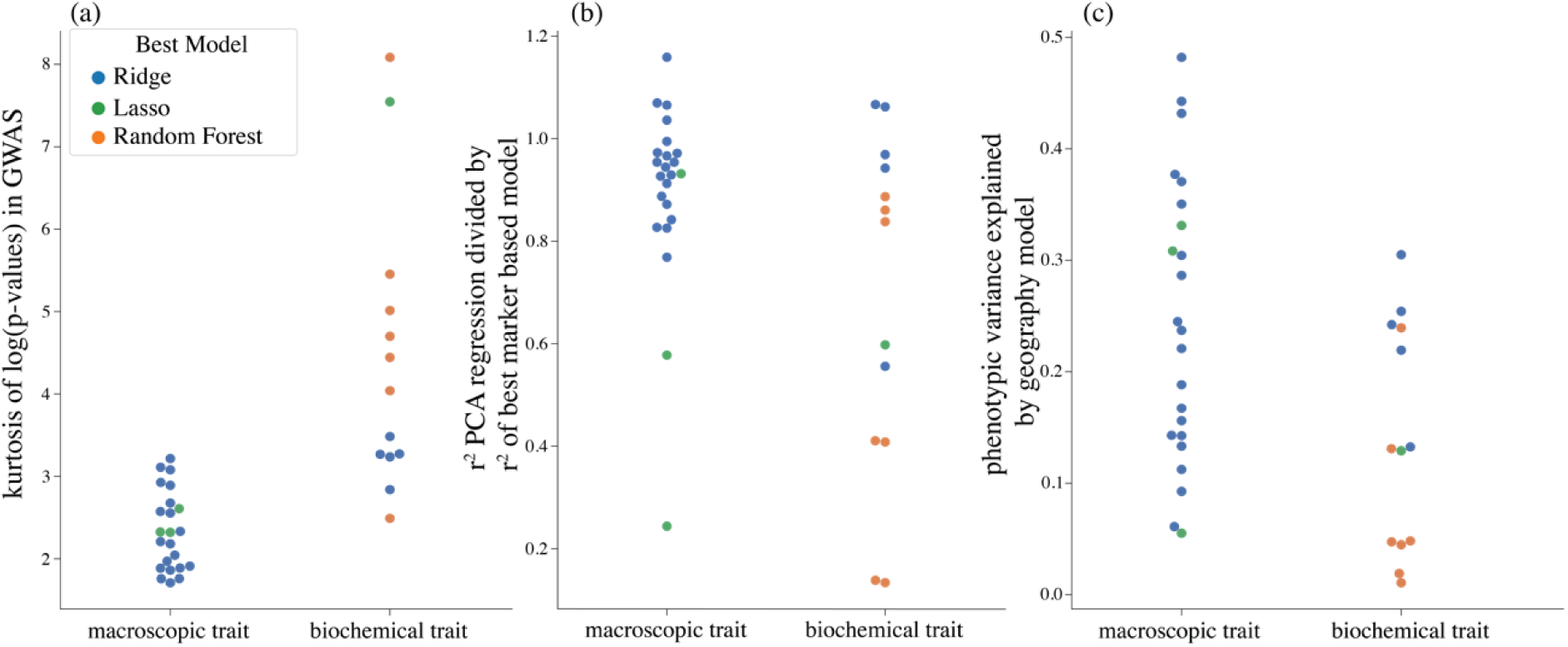
Sensitivity of best genomic prediction model to genetic architecture, population structure and geography. (a) The kurtosis of GWAS log(p-values) for each of the 36 traits analysed in this study. (b) the ratio between prediction accuracy (*r*^2^) of the best performing model fit to the principal components of the genome (30% variance explained), to the best model fit to genome-wide markers. (c) The proportion of phenotypic variance explained by the geographic location of origin of genotypes (*r*^2^), given by equation 2 for each class of trait.

### 3.3 The Effect of genetic architecture and population structure differ across trait ontology

To better understand how population structure affects prediction accuracy, we used principal components (PCs) computed from genome-wide marker data as predictors instead of the markers themselves. We considered traits that could be well predicted by a small number of PCs to be structured. As a quantitative metric of structure we divided the *r*^2^ prediction accuracy of the best model that used markers by the *r*^2^ of the best model using PCs. Under this metric, macroscopic traits were more structured than biochemical traits. When we fitted each model to the PCs explaining 30% of marker variance, the best genomic prediction model fitted to these PCs produced an *r*^2^ value on average 90% as high as the best marker-based model for macroscopic traits, compared to 69% for biochemical traits (t-test, t=2.48, p=0.0178; Fig. 3b).

**Figure 3:**
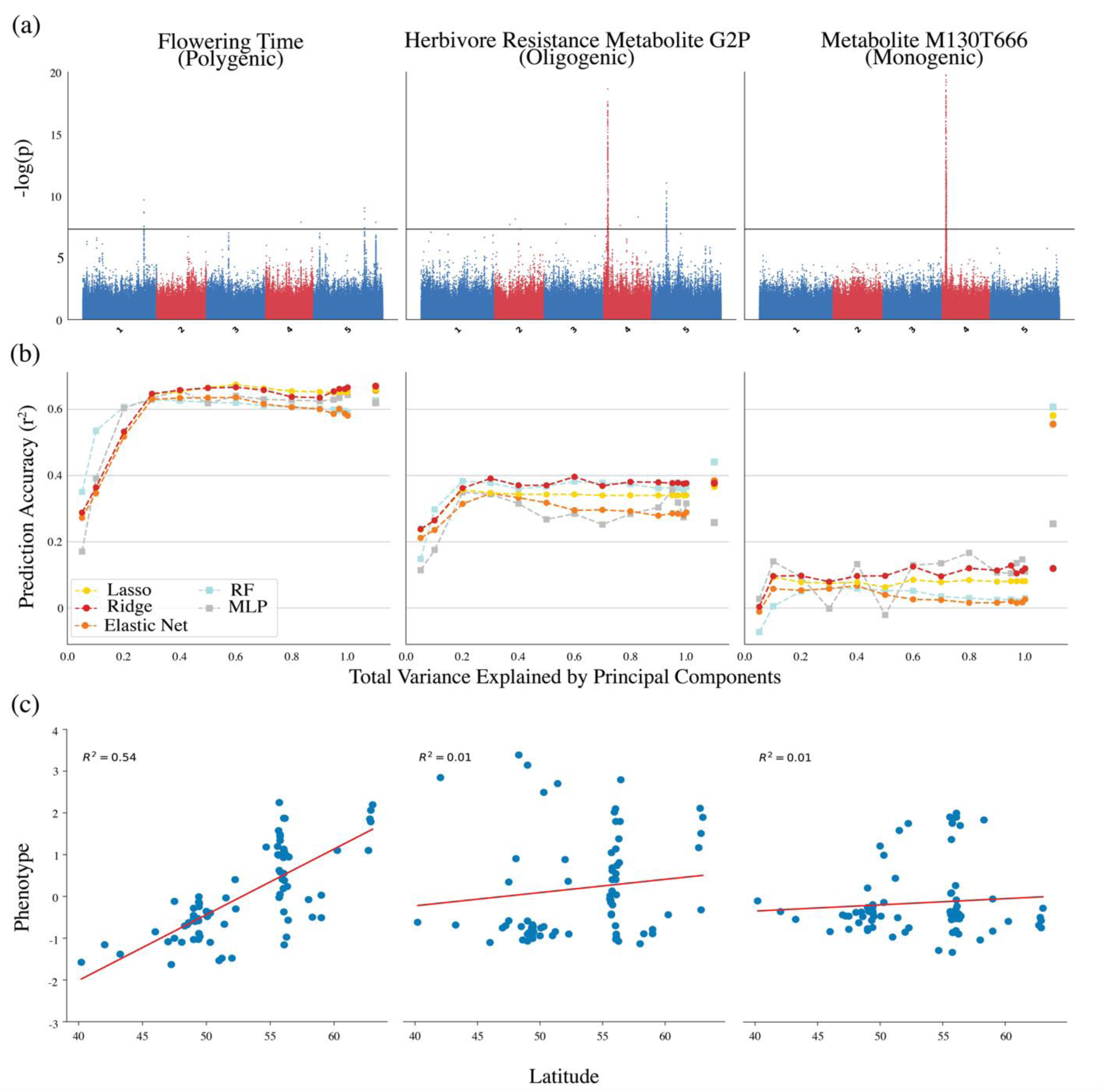
Example of the relationship between genetic architecture, genomic prediction model, population structure and geography. (a) Manhattan plots for GWAS illustrating the genetic architecture of 3 traits with contrasted genetic architectures. (b) Sensitivity to dimensionality reduction conducted through principal component (PC) analysis. The x-axis shows the cumulative variance explained by the set of PCs retained in the untransformed feature set. The last tick on the x-axis corresponds to the accuracy achieved using the markers, equivalent to using all PCs. RF: Random Forest; MLP: multilayer perceptron model which uses a single hidden layer. (c) Sensitivity between the phenotype and the latitude of origin of the accessions. *Note* figure shows only the 78 accessions with phenotype data for all 3 traits.

Finally, we investigated the effect of large spatial gradients. In general, macroscopic traits tended to covary more strongly with latitude and longitude than biochemical traits. A quadratic function of latitude and longitude (Eqn. 2) explained on average 25% of phenotypic variance for macroscopic traits compared to 14% of variance in microscopic traits (t-test, t=2.55, p=0.0153). Notable were the extremes: the quadratic spatial model could explain as much as 48% of flowering time traits, yet as little as 1% of variation in biochemical traits (Fig. 3c). Critically, the traits with a distribution of genetic effects with a low kurtosis were best predicted by a Ridge regression model and explained well by population structure and spatial distribution. In turn, the traits with high kurtosis were best predicted by Lasso or Random Forest and explained poorly by population structure and geography (Fig. 2a-c).

## 4. Discussion

Genomic prediction is an expanding field of research initially motivated by the opportunity of conducting molecular breeding in animals and plants. Genomic selection can apply to any heritable trait, with very diverse genetic architectures and evolutionary histories. As such, we expected that genomic prediction would rely on different, specific methodologies for each trait. Among the 36 *Arabidopsis thaliana* traits tested, the most accurate genomic prediction model depended on trait ontology, genetic architecture and population structure. These components are interconnected and collectively determine the most suitable prediction model. Specifically, we observed that biochemical traits have sparser genetic architectures and are less influenced by population structure. As a result, different approaches were optimal for biochemical and macroscopic traits. Linear regression was best suited for macroscopic traits, while sparse linear regression or Random Forest generally yielded more accurate predictions for biochemical traits.

### Model selection depends on the genetic architecture of the trait

We found that Random Forest specifically outperformed linear models for biochemical traits which showed a monogenic to oligogenic architecture. These findings challenge previous work which has argued that for polygenic traits determined by a complex network of genes, non-linear models have potential to outperform linear models (Azodi et al., 2019; Farooq et al., 2023; Momen et al., 2018; Morgante et al., 2018). Instead, we support that non-linear models are best suited to traits with a small number of effects with strong epistasis (Abdollahi-Arpanahi et al., 2020). Our results also suggest that for training panels gathering broad sources of natural variation, non-linear, non-parametric models can improve predictions when targeted to specific sparse traits. We argue this is a more effective strategy than developing generalised machine learning frameworks aimed at improving genomic prediction across all traits. Our findings are consistent with recent studies that linked the success of ensemble approaches to evidence of strong epistasis in specific mice traits (Perez et al., 2022; Zingaretti et al., 2020), or a recent study finding non-linear approaches to improve prediction of MHC related human disease (Ohta et al., 2024). Our results also expand on a study by Riedelsheimer and colleagues (2012), which compared linear models between molecular and macroscopic traits in maize. Their study found that molecular traits have a simple genetic architecture, and showed that linear models with a sparse prior best predicted them. Consistent with these findings, we generally observed that for biochemical traits the best performing linear models had sparse priors, although random forest delivered higher accuracy for a few of the biochemical traits. Earlier motivation for testing machine learning in complex traits was to capture phenotypic variance underpinned by epistasis. We believe non-parametric models worked specifically well for sparser traits because the presence of interaction effects is stronger when fewer genomic regions underpin a trait. Moreover, when a trait is sparse, the prior distribution of effect sizes is more difficult to determine, which is advantageous to non-parametric models such as Random Forest which rely on fewer assumptions about the prior distribution of genetic effects. The development and regulation of complex traits may well be underpinned by epistasis, but the effect size of each interaction is likely to be small, or underpinned by very rare variants. As noticed in previous studies, this would make the effect size of each interaction small, and difficult to learn from data (Azodi et al., 2019).

In line with previous works, we found artificial neural networks of any design tested to have lower out of sample prediction accuracy than linear models (Abdollahi-Arpanahi et al., 2020; Azodi et al., 2020; Bellot et al., 2018). MLPs make predictions by building increasingly abstract representations from all the predictors. They have the implicit assumption that every predictor is meaningful, and likely interacts with other predictors. These assumptions work well for computer vision tasks, where each pixel contains a small but meaningful amount of information, and an image is best understood from a collection of many pixels. We believe the poor results from MLP in genomic prediction is because the way genomic information maps to phenotypic variation poorly matches the assumptions of MLPs. Genomic data typically exhibit a very low signal-to-noise ratio with respect to trait determinism. For simpler traits, many SNPs may have no signal, and of those that do, there may only be a small number of effective interactions. For complex traits, that are hypothesised to be underpinned by a complex network of genes (Carlborg & Haley, 2004), complex models like MLPs suffer from the aforementioned problem of low statistical power and rare variants (Azodi et al., 2019; Bellot et al., 2018).

### Effect of the trait evolutionary history on genomic prediction

We found that population demography and the evolutionary history of the trait is also an important determinant of genomic prediction model selection. Particularly, biochemical traits were less correlated with population structure than the macroscopic traits. The principal component analysis of the population genomic data captures the major axis of genomic variation, and thus provides a concise summary of population structure (Ma & Amos, 2012; Novembre & Stephens, 2008). For example, a linear combination of PCs explaining 30% of genomic variants predicted flowering time as well as any marker based model. On the other hand, the most accurate model that used PCs as predictors only predicted the concentration of metabolite M130T666 with an *r*^2^ = 22% as high as the best marker based model. Thus in the training population examined, individual markers are informative for the prediction of simple, typically biochemical, traits while for complex, macroscopic traits, causal variants are confounded by population structure. Indeed a complex trait like flowering time may be underpinned by many epistatic effects, but could be obfuscated by population structure, favouring simpler linear models. We tested a population of natural, unrelated accessions sampled at a large geographic scale, which is different to most genomic prediction studies working with inbred lines generated from crossing designs. Genomic prediction in genetically diverse training populations is relevant at the prebreeding step where the aim is often to characterise germplasm as broadly as possible. As a consequence, the effect of population structure we observed might be less pronounced in study with crossing designs. However, simulation studies using genotypic data from crosses have also shown that factoring the effect of populations structure can improve the prediction accuracy of complex traits if the testing partition of the data captures similar variation to the training partition (Fodor et al., 2014).

Our findings suggest that the varying degree of correlation between traits and population structure observed in this study is explained by heterogeneous levels of ancestral selection across traits. We observed that macroscopic traits on average had stronger biological clines across longitudinal and latitudinal gradients compared to biochemical traits. We thus expect that such biological clines are established in response to selection pressures that differ between environments. It is well established that flowering time is adaptive to specific environments in Europe (Fournier-Level et al., 2022). The weaker clines in biochemical traits supports the neutral theory of molecular evolution, which posits that selection occurs mostly at the organismal level and that molecular traits are generally more neutral than macroscopic traits (Fig. 2c; Kimura, 1985). When considering the hierarchical organisation of traits that establish the whole phenotype, it follows that a mutation affecting a trait at a high level will also influence traits at lower levels. Macroscopic traits which are under direct selection combine many endophenotypes. However, individual biochemical traits might not affect the organism at a macroscopic level and thus are inherently under more relaxed selection (Zhang, 2018). We argue that in addition to genetic architecture, ensemble approaches were effective for specifically biochemical traits because these traits are more neutral, and thus less confounded by population structure. Importantly, our research suggests that within the same population, ancestral patterns of selection affect the inference of causal variants and the choice of genomic prediction model. As a consequence, the magnitude of this effect in a natural population can be understood in terms of the hierarchical organisation of trait ontology.

### Conclusion

In light of our results, we suggest that future implementation of genomic prediction should lean towards ensemble models in addition to linear statistical models when the genetic architecture is sparse. We also caution that the confounding effect of population structure can vary between traits within the same population, likely due to selection history. This has the potential to hamper the generalisation and biological interpretability of genomic prediction models. Non-linear approaches have a role in genomic prediction, however only for specific genetic architectures, with strong associations that are not confounded by population structure, or after strong differential selection in the ancestral populations. Previous work has focused on benchmarking a wide range of genomic prediction models across select complex traits without focusing on their organisational hierarchy. Future work would certainly benefit from expanding genomic prediction methodology through the exploration of the phenome, linking trait ontologies, genetic architectures, and population histories to modelling strategies.

## Acknowledgements

We would like to acknowledge Prof. Doug Speed and Prof. David Balding for providing feedback and comments on earlier versions of this manuscript. We acknowledge the excellent support of the Research Computing Services of the University of Melbourne.

## Statements & declarations

### Author contributions

PMG, JFP and AFL designed the study, PMG and JFP prepared the material, PMG analysed the data, PMG and AFL wrote the manuscript.

## References

Abdollahi-Arpanahi, R., Gianola, D., & Peñagaricano, F. (2020). Deep learning versus parametric and ensemble methods for genomic prediction of complex phenotypes. Genetics Selection Evolution, 52(1), 12. 10.1186/s12711-020-00531-z

Alonso-Blanco, C., Andrade, J., Becker, C., Bemm, F., Bergelson, J., Borgwardt, K. M., Cao, J., Chae, E., Dezwaan, T. M., Ding, W., & others. (2016). 1,135 genomes reveal the global pattern of polymorphism in Arabidopsis thaliana. Cell, 166(2), 481–491.

Arouisse, B., Korte, A., van Eeuwijk, F., & Kruijer, W. (2020). Imputation of 3 million SNPs in the Arabidopsis regional mapping population. The Plant Journal: For Cell and Molecular Biology, 102(4), 872–882. 10.1111/tpj.14659

Atwell, S., Huang, Y. S., Vilhjálmsson, B. J., Willems, G., Horton, M., Li, Y., Meng, D., Platt, A., Tarone, A. M., Hu, T. T., Jiang, R., Muliyati, N. W., Zhang, X., Amer, M. A., Baxter, I., Brachi, B., Chory, J., Dean, C., Debieu, M., … Nordborg, M. (2010). Genome-wide association study of 107 phenotypes in Arabidopsis thaliana inbred lines. Nature, 465(7298), 627–631. 10.1038/nature08800

Azodi, C. B., Bolger, E., McCarren, A., Roantree, M., de los Campos, G., & Shiu, S.-H. (2019). Benchmarking Parametric and Machine Learning Models for Genomic Prediction of Complex Traits. G3 Genes|Genomes|Genetics, 9(11), 3691–3702. 10.1534/g3.119.400498

Azodi, C. B., Pardo, J., VanBuren, R., de los Campos, G., & Shiu, S.-H. (2020). Transcriptome-Based Prediction of Complex Traits in Maize[OPEN]. The Plant Cell, 32(1), 139–151. 10.1105/tpc.19.00332

Bellot, P., de los Campos, G., & Pérez-Enciso, M. (2018). Can Deep Learning Improve Genomic Prediction of Complex Human Traits? Genetics, 210(3), 809–819. 10.1534/genetics.118.301298

Brachi, B., Meyer, C. G., Villoutreix, R., Platt, A., Morton, T. C., Roux, F., & Bergelson, J. (2015). Coselected genes determine adaptive variation in herbivore resistance throughout the native range of Arabidopsis thaliana. Proceedings of the National Academy of Sciences, 112(13), 4032–4037. 10.1073/pnas.1421416112

Carlborg, Ö., & Haley, C. S. (2004). Epistasis: Too often neglected in complex trait studies? Nature Reviews Genetics, 5(8), 618–625. 10.1038/nrg1407

Cooper, L., Elser, J., Laporte, M.-A., Arnaud, E., & Jaiswal, P. (2024). Planteome 2024 Update: Reference Ontologies and Knowledgebase for Plant Biology. Nucleic Acids Research, 52(D1), D1548–D1555. 10.1093/nar/gkad1028

Farooq M., van Dijk A.D.J,. Nijveen H., Mansoor S., de Ridder D. (2022). Genomic prediction in plants: opportunities for ensemble machine learning based approaches. F1000 Research., 18;11:802. 10.12688/f1000research.122437.2

Fodor, A., Segura, V., Denis, M., Neuenschwander, S., Fournier-Level, A., Chatelet, P., Homa, F. A. A., Lacombe, T., This, P., & Le Cunff, L. (2014). Genome-Wide Prediction Methods in Highly Diverse and Heterozygous Species: Proof-of-Concept through Simulation in Grapevine. PLOS ONE, 9(11), 1–14. 10.1371/journal.pone.0110436

Fournier-Level, A., Taylor, M. A., Paril, J. F., Martínez-Berdeja, A., Stitzer, M. C., Cooper, M. D., Roe, J. L., Wilczek, A. M., & Schmitt, J. (2022). Adaptive significance of flowering time variation across natural seasonal environments in Arabidopsis thaliana. New Phytologist, 234(2), 719–734. 10.1111/nph.17999

Guo, Z., Tucker, D. M., Basten, C. J., Gandhi, H., Ersoz, E., Guo, B., Xu, Z., Wang, D., & Gay, G. (2014). The impact of population structure on genomic prediction in stratified populations. Theoretical and Applied Genetics, 127(3), 749–762. 10.1007/s00122-013-2255-x

Horton, M. W., Hancock, A. M., Huang, Y. S., Toomajian, C., Atwell, S., Auton, A., Muliyati, N. W., Platt, A., Sperone, F. G., Vilhjálmsson, B. J., Nordborg, M., Borevitz, J. O., & Bergelson, J. (2012). Genome-wide patterns of genetic variation in worldwide Arabidopsis thaliana accessions from the RegMap panel. Nature Genetics, 44(2), 212–216. 10.1038/ng.1042

Jumper, J., Evans, R., Pritzel, A., Green, T., Figurnov, M., Ronneberger, O., Tunyasuvunakool, K., Bates, R., Žídek, A., Potapenko, A., & others. (2021). Highly accurate protein structure prediction with AlphaFold. Nature, 596(7873), 583–589.

Kimura, M. (1985). The neutral theory of molecular evolution. CambRidge University Press.

Kingma, D. P., & Ba, J. (2014). Adam: A method for stochastic optimization. arXiv Preprint arXiv:1412.6980.

Ma, J., & Amos, C. I. (2012). Principal components analysis of population admixture. PloS One, 7(7), e40115.

Meuwissen, T. H. (2009). Accuracy of breeding values of “unrelated” individuals predicted by dense SNP genotyping. Genetics Selection Evolution, 41(1), 35. 10.1186/1297-9686-41-35

Meuwissen, T. H., Hayes, B. J., & Goddard, M. E. (2001). Prediction of Total Genetic Value Using Genome-Wide Dense Marker Maps. Genetics, 157(4), 1819–1829. 10.1093/genetics/157.4.1819

Momen, M., Mehrgardi, A. A., Sheikhi, A., Kranis, A., Tusell, L., Morota, G., Rosa, G. J. M., & Gianola, D. (2018). Predictive ability of genome-assisted statistical models under various forms of gene action. Scientific Reports, 8(1), 12309. 10.1038/s41598-018-30089-2

Morgante, F., Huang, W., Maltecca, C., & Mackay, T. F. C. (2018). Effect of genetic architecture on the prediction accuracy of quantitative traits in samples of unrelated individuals. Heredity, 120(6), 500–514. 10.1038/s41437-017-0043-0

Novembre, J., & Stephens, M. (2008). Interpreting principal component analyses of spatial population genetic variation. Nature Genetics, 40(5), 646–649.

Ohta, R., Tanigawa, Y., Suzuki, Y., Kellis, M., & Morishita, S. (2024). A polygenic score method boosted by non-additive models. Nature Communications, 15(1), 4433.

Pedregosa, F., Varoquaux, G., Gramfort, A., Michel, V., Thirion, B., Grisel, O., Blondel, M., Prettenhofer, P., Weiss, R., Dubourg, V., Vanderplas, J., Passos, A., Cournapeau, D., Brucher, M., Perrot, M., & Duchesnay, E. (2011). Scikit-learn: Machine Learning in Python. Journal of Machine Learning Research, 12, 2825–2830.

Perez, B. C., Bink, M. C. A. M., Svenson, K. L., Churchill, G. A., & Calus, M. P. L. (2022). Prediction performance of linear models and gradient boosting machine on complex phenotypes in outbred mice. G3 Genes|Genomes|Genetics, 12(4), jkac039. 10.1093/g3journal/jkac039

Prechelt, L. (1998). Automatic early stopping using cross validation: Quantifying the criteria. Neural Networks, 11(4), 761–767.

Putra, A. R., Yen, J. D. L., & Fournier-Level, A. (2023). Forecasting trait responses in novel environments to aid seed provenancing under climate change. Molecular Ecology Resources, 23(3), 565–580. 10.1111/1755-0998.13728

Riedelsheimer, C., Technow, F., & Melchinger, A. E. (2012). Comparison of whole-genome prediction models for traits with contrasting genetic architecture in a diversity panel of maize inbred lines. BMC Genomics, 13(1), 452. 10.1186/1471-2164-13-452

Seren, Ü., Grimm, D., Fitz, J., Weigel, D., Nordborg, M., Borgwardt, K., & Korte, A. (2017). AraPheno: A public database for Arabidopsis thaliana phenotypes. Nucleic Acids Research, 45(D1), D1054– D1059. 10.1093/nar/gkw986

Šimić, D., Mladenović Drinić, S., Zdunić, Z., Jambrović, A., Ledenčan, T., Brkić, J., Brkić, A., & Brkić, I. (2012). Quantitative trait loci for biofortification traits in maize grain. Journal of Heredity, 103(1), 47–54.

Srivastava, N., Hinton, G., Krizhevsky, A., Sutskever, I., & Salakhutdinov, R. (2014). Dropout: A simple way to prevent neural networks from overfitting. The Journal of Machine Learning Research, 15(1), 1929–1958.

Tibshirani, R. (1996). Regression Shrinkage and Selection Via the Lasso. Journal of the Royal Statistical Society: Series B (Methodological), 58(1), 267–288. 10.1111/j.2517-6161.1996.tb02080.x

Vogt, F., Shirsekar, G., & Weigel, D. (2022). vcf2gwas: Python API for comprehensive GWAS analysis using GEMMA. Bioinformatics, 38(3), 839–840. 10.1093/bioinformatics/btab710

Zan, Y., & Carlborg, Ö. (2019). A Polygenic Genetic Architecture of Flowering Time in the Worldwide Arabidopsis thaliana Population. Molecular Biology and Evolution, 36(1), 141–154. 10.1093/molbev/msy203

Zhang, J. (2018). Neutral theory and phenotypic evolution. Molecular Biology and Evolution, 35(6), 1327–1331.

Zhou, X., & Stephens, M. (2012). Genome-wide efficient mixed-model analysis for association studies. Nature Genetics, 44(7), 821–824. 10.1038/ng.2310

Zingaretti, L. M., Gezan, S. A., Ferrão, L. F. V., Osorio, L. F., Monfort, A., Muñoz, P. R., Whitaker, V. M., & Pérez-Enciso, M. (2020). Exploring deep learning for complex trait genomic prediction in polyploid outcrossing species. Frontiers in Plant Science, 11, 25.

Zou, H., & Hastie, T. (2005). Regularization and variable selection via the Elastic-net. Journal of the Royal Statistical Society Series B: Statistical Methodology, 67(2), 301–320.

