## Supplementary Materials for "Trait genetic architecture and population structure determine model selection for genomic prediction in natural *Arabidopsis thaliana* populations"

### **Supplementary Table S1**: *Arabidopsis thaliana* traits analysed

The following table summarises the traits selected from all of the traits available in the AraPheno database, as well as those measured by Brachi and colleagues (2015).

| **Name Abbreviation** | **description** | **Arapheno Study Number** | **Trait ontology** | **Number of accessions** | **Citation** |
| --- | --- | --- | --- | --- | --- |
| G2H3B | Concentration of the methionine-derived Glucosinolate G2H3B in dried rosette tissue | Not listed on Arapheno | biochemical trait (TO:0000277) | 595 | Brachi, B., Meyer, C. G., Villoutreix, R., Platt, A., Morton, T. C., Roux, F., & Bergelson, J. (2015). Coselected genes determine adaptive variation in herbivore resistance throughout the native range of Arabidopsis thaliana. *Proceedings of the National Academy of Sciences*, *112*(13), 4032-4037 |
| G2H4P | Concentration of the methionine-derived Glucosinolate G2H4P in dried rosette tissue | Not listed on Arapheno | biochemical trait (TO:0000277) | 595 | Brachi, B., Meyer, C. G., Villoutreix, R., Platt, A., Morton, T. C., Roux, F., & Bergelson, J. (2015). Coselected genes determine adaptive variation in herbivore resistance throughout the native range of Arabidopsis thaliana. *Proceedings of the National Academy of Sciences*, *112*(13), 4032-4037 |
| G2P | Concentration of the methionine-derived Glucosinolate G2P in dried rosette tissue | Not listed on Arapheno | biochemical trait (TO:0000277) | 595 | Brachi, B., Meyer, C. G., Villoutreix, R., Platt, A., Morton, T. C., Roux, F., & Bergelson, J. (2015). Coselected genes determine adaptive variation in herbivore resistance throughout the native range of Arabidopsis thaliana. *Proceedings of the National Academy of Sciences*, *112*(13), 4032-4037 |
| G3B | Concentration of the methionine-derived Glucosinolate G3B in dried rosette tissue | Not listed on Arapheno | biochemical trait (TO:0000277) | 595 | Brachi, B., Meyer, C. G., Villoutreix, R., Platt, A., Morton, T. C., Roux, F., & Bergelson, J. (2015). Coselected genes determine adaptive variation in herbivore resistance throughout the native range of Arabidopsis thaliana. *Proceedings of the National Academy of Sciences*, *112*(13), 4032-4037 |
| G3HP | Concentration of the methionine-derived Glucosinolate G3HP in dried rosette tissue | Not listed on Arapheno | biochemical trait (TO:0000277) | 595 | Brachi, B., Meyer, C. G., Villoutreix, R., Platt, A., Morton, T. C., Roux, F., & Bergelson, J. (2015). Coselected genes determine adaptive variation in herbivore resistance throughout the native range of Arabidopsis thaliana. *Proceedings of the National Academy of Sciences*, *112*(13), 4032-4037 |
| G4MSB | Concentration of the methionine-derived Glucosinolate G4MSB in dried rosette tissue | Not listed on Arapheno | biochemical trait (TO:0000277) | 595 | Brachi, B., Meyer, C. G., Villoutreix, R., Platt, A., Morton, T. C., Roux, F., & Bergelson, J. (2015). Coselected genes determine adaptive variation in herbivore resistance throughout the native range of Arabidopsis thaliana. *Proceedings of the National Academy of Sciences*, *112*(13), 4032-4037 |
| G4P | Concentration of the methionine-derived Glucosinolate G4P in dried rosette tissue | Not listed on Arapheno | biochemical trait (TO:0000277) | 595 | Brachi, B., Meyer, C. G., Villoutreix, R., Platt, A., Morton, T. C., Roux, F., & Bergelson, J. (2015). Coselected genes determine adaptive variation in herbivore resistance throughout the native range of Arabidopsis thaliana. *Proceedings of the National Academy of Sciences*, *112*(13), 4032-4037 |
| G5MSP | Concentration of the methionine-derived Glucosinolate G5MSP in dried rosette tissue | Not listed on Arapheno | biochemical trait (TO:0000277) | 595 | Brachi, B., Meyer, C. G., Villoutreix, R., Platt, A., Morton, T. C., Roux, F., & Bergelson, J. (2015). Coselected genes determine adaptive variation in herbivore resistance throughout the native range of Arabidopsis thaliana. *Proceedings of the National Academy of Sciences*, *112*(13), 4032-4037 |
| Cd111 | Cadmium concentration in leaves. Plants grown in a greenhouse environment. After 5 weeks plants were non-destructively sampled by removing one or two leaves and the elemental composition of the tissue analyzed by Inductively Couple Plasma Mass Spectroscopy (ICP-MS). | 16 | biochemical trait (TO:0000277) | 345 | Baxter, I., Brazelton, J. N., Yu, D., Huang, Y. S., Lahner, B., Yakubova, E., ... & Salt, D. E. (2010). A coastal cline in sodium accumulation in Arabidopsis thaliana is driven by natural variation of the sodium transporter AtHKT1; 1. *PLoS genetics*, *6*(11), e1001193. |
| Mo98 | Molybdenum concentration in leaves. Plants grown in a greenhouse environment. After 5 weeks plants were non-destructively sampled by removing one or two leaves and the elemental composition of the tissue analyzed by Inductively Couple Plasma Mass Spectroscopy (ICP-MS). | 16 | biochemical trait (TO:0000277) | 345 | Baxter, I., Brazelton, J. N., Yu, D., Huang, Y. S., Lahner, B., Yakubova, E., ... & Salt, D. E. (2010). A coastal cline in sodium accumulation in Arabidopsis thaliana is driven by natural variation of the sodium transporter AtHKT1; 1. *PLoS genetics*, *6*(11), e1001193. |
| FLC | Gene expression of Flowering Locus C gene. RNA was extracted from leaves after 4 wks of growth. FLC gene expression levels were determined by Northern hybridization quantified relative to Beta;-TUBULIN expression | 1 | biochemical trait (TO:0000277) | 167 | Atwell S, Huang YS, Vilhjálmsson BJ, Willems G, Horton M, Li Y, Meng D, Platt A, Tarone AM, Hu TT, Jiang R, Muliyati NW, Zhang X, Amer MA, Baxter I, Brachi B, Chory J, Dean C, Debieu M, de Meaux J, Ecker JR, Faure N, Kniskern JM, Jones JD, Michael T, Nemri A, Roux F, Salt DE, Tang C, Todesco M, Traw MB, Weigel D, Marjoram P, Borevitz JO, Bergelson J, Nordborg M. (2010).  Genome-wide association study of 107 phenotypes in Arabidopsis thaliana inbred lines.  Nature. 465(7298). 627-31.  doi:10.1038/nature08800 |
| FRI | Gene Expression of frigida. RNA was extracted from leaves after 4 wks of growth. FRI gene expression levels were determined by Northern hybridization quantified relative to Beta;-TUBULIN expression | 1 | biochemical trait (TO:0000277) | 164 | Atwell S, Huang YS, Vilhjálmsson BJ, Willems G, Horton M, Li Y, Meng D, Platt A, Tarone AM, Hu TT, Jiang R, Muliyati NW, Zhang X, Amer MA, Baxter I, Brachi B, Chory J, Dean C, Debieu M, de Meaux J, Ecker JR, Faure N, Kniskern JM, Jones JD, Michael T, Nemri A, Roux F, Salt DE, Tang C, Todesco M, Traw MB, Weigel D, Marjoram P, Borevitz JO, Bergelson J, Nordborg M. (2010).  Genome-wide association study of 107 phenotypes in Arabidopsis thaliana inbred lines.  Nature. 465(7298). 627-31.  doi:10.1038/nature08800 |
| M130T666 | LC-MS–based untargeted metabolite profiles of leaf tissue were collected for each accession. Metabolite feature with a mass-to-charge ratio of 130 and a retention time of 666s | 4 | biochemical trait (TO:0000277) | 405 | Strauch, R. C., Svedin, E., Dilkes, B., Chapple, C., & Li, X. (2015). Discovery of a novel amino acid racemase through exploration of natural variation in Arabidopsis thaliana. *Proceedings of the National Academy of Sciences*, *112*(37), 11726-11731. |
| FT10 | days to flowering trait  In a greenhouse at 10 degrees. Flowering time was scored as days until first  open flower | 12 | flowering time trait (TO:0002616) | 1058 | Alonso-Blanco, C., Andrade, J., Becker, C., Bemm, F., Bergelson, J., Borgwardt, K. M., ... & Zhou, X. (2016). 1,135 genomes reveal the global pattern of polymorphism in Arabidopsis thaliana. *Cell*, *166*(2), 481-491. |
| 0W | Flowering time trait. Number of days required for bolt height to reach 5cm | 1 | flowering time trait (TO:0002616) | 137 | Atwell S, Huang YS, Vilhjálmsson BJ, Willems G, Horton M, Li Y, Meng D, Platt A, Tarone AM, Hu TT, Jiang R, Muliyati NW, Zhang X, Amer MA, Baxter I, Brachi B, Chory J, Dean C, Debieu M, de Meaux J, Ecker JR, Faure N, Kniskern JM, Jones JD, Michael T, Nemri A, Roux F, Salt DE, Tang C, Todesco M, Traw MB, Weigel D, Marjoram P, Borevitz JO, Bergelson J, Nordborg M. (2010).  Genome-wide association study of 107 phenotypes in Arabidopsis thaliana inbred lines.  Nature. 465(7298). 627-31.  doi:10.1038/nature08800 |
| 8W_GH_FT | Flowering time trait. Number of days required for bolt height to reach 5cm | 1 | flowering time trait (TO:0002616) | 162 | Atwell S, Huang YS, Vilhjálmsson BJ, Willems G, Horton M, Li Y, Meng D, Platt A, Tarone AM, Hu TT, Jiang R, Muliyati NW, Zhang X, Amer MA, Baxter I, Brachi B, Chory J, Dean C, Debieu M, de Meaux J, Ecker JR, Faure N, Kniskern JM, Jones JD, Michael T, Nemri A, Roux F, Salt DE, Tang C, Todesco M, Traw MB, Weigel D, Marjoram P, Borevitz JO, Bergelson J, Nordborg M. (2010).  Genome-wide association study of 107 phenotypes in Arabidopsis thaliana inbred lines.  Nature. 465(7298). 627-31.  doi:10.1038/nature08800 |
| FT22 | days to emergence of first buds in the greenhouse at 22 degrees. Plants were checked bi-weekly for presence of first buds, and the average flowering time of 4 plants of the same accession were collected | 1 | flowering time trait (TO:0002616) | 193 | Atwell S, Huang YS, Vilhjálmsson BJ, Willems G, Horton M, Li Y, Meng D, Platt A, Tarone AM, Hu TT, Jiang R, Muliyati NW, Zhang X, Amer MA, Baxter I, Brachi B, Chory J, Dean C, Debieu M, de Meaux J, Ecker JR, Faure N, Kniskern JM, Jones JD, Michael T, Nemri A, Roux F, Salt DE, Tang C, Todesco M, Traw MB, Weigel D, Marjoram P, Borevitz JO, Bergelson J, Nordborg M. (2010).  Genome-wide association study of 107 phenotypes in Arabidopsis thaliana inbred lines.  Nature. 465(7298). 627-31.  doi:10.1038/nature08800 |
| FT_Field | Flowering time measured in a common field experiment. Flowering time was scored as the number of days between germination date and appearance of the first flower | NaN | flowering time trait (TO:0002616) | 180 | Atwell S, Huang YS, Vilhjálmsson BJ, Willems G, Horton M, Li Y, Meng D, Platt A, Tarone AM, Hu TT, Jiang R, Muliyati NW, Zhang X, Amer MA, Baxter I, Brachi B, Chory J, Dean C, Debieu M, de Meaux J, Ecker JR, Faure N, Kniskern JM, Jones JD, Michael T, Nemri A, Roux F, Salt DE, Tang C, Todesco M, Traw MB, Weigel D, Marjoram P, Borevitz JO, Bergelson J, Nordborg M. (2010).  Genome-wide association study of 107 phenotypes in Arabidopsis thaliana inbred lines.  Nature. 465(7298). 627-31.  doi:10.1038/nature08800 |
| SDV | Number of days following stratification to opening of first flower. The experiment was stopped at 200 d, and accessions that had not flowered at that point were assigned a value of 200 | 1 | flowering time trait (TO:0002616) | 159 | Atwell S, Huang YS, Vilhjálmsson BJ, Willems G, Horton M, Li Y, Meng D, Platt A, Tarone AM, Hu TT, Jiang R, Muliyati NW, Zhang X, Amer MA, Baxter I, Brachi B, Chory J, Dean C, Debieu M, de Meaux J, Ecker JR, Faure N, Kniskern JM, Jones JD, Michael T, Nemri A, Roux F, Salt DE, Tang C, Todesco M, Traw MB, Weigel D, Marjoram P, Borevitz JO, Bergelson J, Nordborg M. (2010).  Genome-wide association study of 107 phenotypes in Arabidopsis thaliana inbred lines.  Nature. 465(7298). 627-31.  doi:10.1038/nature08800 |
| CL | Cauline Leaf Number. Seeds for 1135 Arabidopsis accessions (1001 Genomes Consortium, 2016) were surface-sterilized in 95% ethanol for 5 min and allowed to air- dry. After 6 d of stratification in the dark at 4°C in 0.1% agarose, seeds were distributed across 4800 pots as four replicates in a randomized block design, with each replicate corresponding to one block. Plants were grown in controlled growth chambers with the following settings: 16 h light/8 h darkness, 16°C constant temperature, 65% humidity. All trays within a block were moved to a new shelf and rotated 180°C every other day to minimize position effects. | 38 | plant [organ morphology trait](https://bioportal.bioontology.org/ontologies/PTO/?p=classes&conceptid=http%3A%2F%2Fpurl.obolibrary.org%2Fobo%2FTO_0000736) (TO:0000836) | 904 | Alonso-Blanco, C., Andrade, J., Becker, C., Bemm, F., Bergelson, J., Borgwardt, K. M., ... & Zhou, X. (2016). 1,135 genomes reveal the global pattern of polymorphism in Arabidopsis thaliana. *Cell*, *166*(2), 481-491. |
| RL | Rosette Leaf Number. Seeds for 1135 Arabidopsis accessions (1001 Genomes Consortium, 2016) were surface-sterilized in 95% ethanol for 5 min and allowed to air- dry. After 6 d of stratification in the dark at 4°C in 0.1% agarose, seeds were distributed across 4800 pots as four replicates in a randomized block design, with each replicate corresponding to one block. Plants were grown in controlled growth chambers with the following settings: 16 h light/8 h darkness, 16°C constant temperature, 65% humidity. All trays within a block were moved to a new shelf and rotated 180°C every other day to minimize position effects. | 38 | plant [organ morphology trait](https://bioportal.bioontology.org/ontologies/PTO/?p=classes&conceptid=http%3A%2F%2Fpurl.obolibrary.org%2Fobo%2FTO_0000736) (TO:0000836) | 850 | Alonso-Blanco, C., Andrade, J., Becker, C., Bemm, F., Bergelson, J., Borgwardt, K. M., ... & Zhou, X. (2016). 1,135 genomes reveal the global pattern of polymorphism in Arabidopsis thaliana. *Cell*, *166*(2), 481-491. |
| GR21 | Seed dormancy. Seeds from three biological replicates per line were pooled together to compensate for growth chamber heterogeneity. About 100 seeds per genotype were spread on wet filter paper in Petri dishes subsequently placed in moisture chambers (following protocol of Alonso-Blanco et al., 2003) that were located in a culture room set at 25°C with a long-day light regime (16 hr light, 8 hr dark). Germination rate (GR21) was determined after 7 days of incubation by scoring radicle emergence. | 19 | seed dormancy trait (TO:0000253) | 161 | Togninalli, M., Seren, Ü., Freudenthal, J. A., Monroe, J. G., Meng, D., Nordborg, M., ... & Grimm, D. G. (2020). AraPheno and the AraGWAS Catalog 2020: a major database update including RNA-Seq and knockout mutation data for Arabidopsis thaliana. *Nucleic acids research*, *48*(D1), D1063-D1068. |
| 8W_GH_LN | Leaf number was scored as the number of rosette leaves and the number of cauline leaves when the bolt reached 5cm | 1 | shoot system growth and development trait (TO:0000928) | 163 | Atwell S, Huang YS, Vilhjálmsson BJ, Willems G, Horton M, Li Y, Meng D, Platt A, Tarone AM, Hu TT, Jiang R, Muliyati NW, Zhang X, Amer MA, Baxter I, Brachi B, Chory J, Dean C, Debieu M, de Meaux J, Ecker JR, Faure N, Kniskern JM, Jones JD, Michael T, Nemri A, Roux F, Salt DE, Tang C, Todesco M, Traw MB, Weigel D, Marjoram P, Borevitz JO, Bergelson J, Nordborg M. (2010).  Genome-wide association study of 107 phenotypes in Arabidopsis thaliana inbred lines.  Nature. 465(7298). 627-31.  doi:10.1038/nature08800 |
| LN10 | Leaf number in greenhouse at 10 degrees. Plants were checked bi-weekly for presence of first buds, and the average leaf number at flowering time of 4 plants of the same accession were collected | 1 | shoot system growth and development trait (TO:0000928) | 177 | Atwell S, Huang YS, Vilhjálmsson BJ, Willems G, Horton M, Li Y, Meng D, Platt A, Tarone AM, Hu TT, Jiang R, Muliyati NW, Zhang X, Amer MA, Baxter I, Brachi B, Chory J, Dean C, Debieu M, de Meaux J, Ecker JR, Faure N, Kniskern JM, Jones JD, Michael T, Nemri A, Roux F, Salt DE, Tang C, Todesco M, Traw MB, Weigel D, Marjoram P, Borevitz JO, Bergelson J, Nordborg M. (2010).  Genome-wide association study of 107 phenotypes in Arabidopsis thaliana inbred lines.  Nature. 465(7298). 627-31.  doi:10.1038/nature08800 |
| LN16 | Leaf number in greenhouse at 16 degrees.. Plants were checked bi-weekly for presence of first buds, and the average leaf number at flowering time of 4 plants of the same accession were collected | 1 | shoot system growth and development trait (TO:0000928) | 176 | Atwell S, Huang YS, Vilhjálmsson BJ, Willems G, Horton M, Li Y, Meng D, Platt A, Tarone AM, Hu TT, Jiang R, Muliyati NW, Zhang X, Amer MA, Baxter I, Brachi B, Chory J, Dean C, Debieu M, de Meaux J, Ecker JR, Faure N, Kniskern JM, Jones JD, Michael T, Nemri A, Roux F, Salt DE, Tang C, Todesco M, Traw MB, Weigel D, Marjoram P, Borevitz JO, Bergelson J, Nordborg M. (2010).  Genome-wide association study of 107 phenotypes in Arabidopsis thaliana inbred lines.  Nature. 465(7298). 627-31.  doi:10.1038/nature08800 |
| Storage_28_days | Primary dormancy was measured as the progressive increase of germination rate measured after 28 days of dry storage | NaN | seed dormancy trait (TO:0000253) | 110 | Atwell S, Huang YS, Vilhjálmsson BJ, Willems G, Horton M, Li Y, Meng D, Platt A, Tarone AM, Hu TT, Jiang R, Muliyati NW, Zhang X, Amer MA, Baxter I, Brachi B, Chory J, Dean C, Debieu M, de Meaux J, Ecker JR, Faure N, Kniskern JM, Jones JD, Michael T, Nemri A, Roux F, Salt DE, Tang C, Todesco M, Traw MB, Weigel D, Marjoram P, Borevitz JO, Bergelson J, Nordborg M. (2010).  Genome-wide association study of 107 phenotypes in Arabidopsis thaliana inbred lines.  Nature. 465(7298). 627-31.  doi:10.1038/nature08800 |
| Storage_56_days | Primary dormancy was measured as the progressive increase of germination rate measured after 28 days of dry storage | NaN | seed dormancy trait (TO:0000253) | 110 | Atwell S, Huang YS, Vilhjálmsson BJ, Willems G, Horton M, Li Y, Meng D, Platt A, Tarone AM, Hu TT, Jiang R, Muliyati NW, Zhang X, Amer MA, Baxter I, Brachi B, Chory J, Dean C, Debieu M, de Meaux J, Ecker JR, Faure N, Kniskern JM, Jones JD, Michael T, Nemri A, Roux F, Salt DE, Tang C, Todesco M, Traw MB, Weigel D, Marjoram P, Borevitz JO, Bergelson J, Nordborg M. (2010).  Genome-wide association study of 107 phenotypes in Arabidopsis thaliana inbred lines.  Nature. 465(7298). 627-31.  doi:10.1038/nature08800 |
| Width_10 | The diameters of 4 plants of each accession were measured and the results were expressed as an average value across all available replicates. Plants were scored 8 weeks post germination | NaN | shoot system growth and development trait (TO:0000928) | 176 | Atwell S, Huang YS, Vilhjálmsson BJ, Willems G, Horton M, Li Y, Meng D, Platt A, Tarone AM, Hu TT, Jiang R, Muliyati NW, Zhang X, Amer MA, Baxter I, Brachi B, Chory J, Dean C, Debieu M, de Meaux J, Ecker JR, Faure N, Kniskern JM, Jones JD, Michael T, Nemri A, Roux F, Salt DE, Tang C, Todesco M, Traw MB, Weigel D, Marjoram P, Borevitz JO, Bergelson J, Nordborg M. (2010).  Genome-wide association study of 107 phenotypes in Arabidopsis thaliana inbred lines.  Nature. 465(7298). 627-31.  doi:10.1038/nature08800 |
| MeanTRL_C | total root length. After 3 days of treatment, plates were scanned with CCD flatbed scanners (EPSON Perfection V600 Photo, Seiko Epson, Nagano, Japan), and images used to quantify root parameters with FIJI, by using the tool ‘segmented line’ (Schindelin et al., 2012) as described in Ristova and Busch (2017). In particular, we quantified: primary root length on day 10 (P), growth rate of P after treatment (P2), branching zone or the length of P between the first and last visible lateral root (R), average lateral root length (LRL), and visible lateral root number (LR.No). | 27 | plant [organ morphology trait](https://bioportal.bioontology.org/ontologies/PTO/?p=classes&conceptid=http%3A%2F%2Fpurl.obolibrary.org%2Fobo%2FTO_0000736) (TO:0000836) | 190 | Ristova, D., Giovannetti, M., Metesch, K., & Busch, W. (2018). Natural genetic variation shapes root system responses to phytohormones in Arabidopsis. *The Plant Journal*, *96*(2), 468-481. |
| MeanTRL_CK | Total root length, with cytokine treatment. After 3 days of treatment, plates were scanned with CCD flatbed scanners (EPSON Perfection V600 Photo, Seiko Epson, Nagano, Japan), and images used to quantify root parameters with FIJI, by using the tool ‘segmented line’ (Schindelin et al., 2012) as described in Ristova and Busch (2017). In particular, we quantified: primary root length on day 10 (P), growth rate of P after treatment (P2), branching zone or the length of P between the first and last visible lateral root (R), average lateral root length (LRL), and visible lateral root number (LR.No). | 30 | plant [organ morphology trait](https://bioportal.bioontology.org/ontologies/PTO/?p=classes&conceptid=http%3A%2F%2Fpurl.obolibrary.org%2Fobo%2FTO_0000736) (TO:0000836) | 192 | Ristova, D., Giovannetti, M., Metesch, K., & Busch, W. (2018). Natural genetic variation shapes root system responses to phytohormones in Arabidopsis. *The Plant Journal*, *96*(2), 468-481. |
| DTFmainEffect2009 | days to flowering trait in simulated weather and seasons in greenhouse experiments. Each accession in each experiment had four replicates under each of the four growth conditions (two planting seasons by two locations). Trait effect estimated from two experiments in two walk-in growth chambers (AR-916, Percival Scientific) that were programmed to cycle the local climates every 5 min from the simulated weather files. The simulated climates were generated by using SolarCalc (35) with sunrise and sunset, light spectrum, temperature, and relative humidity programmed to cycle throughout the day and the season according to 1975?2000 averages (Fig. S5). One chamber was simulating Spain (latitude 41.72091, longitude 2.957075) starting from March 1, and the other chamber was simulating Sweden (latitude 55.71226, longitude 13.207352) starting from May 1st on the first day when plants were put into the chambers. | 2 | flowering time trait (TO:0002616) | 468 | Li, Y., Huang, Y., Bergelson, J., Nordborg, M., & Borevitz, J. O. (2010). Association mapping of local climate-sensitive quantitative trait loci in Arabidopsis thaliana. *Proceedings of the National Academy of Sciences*, *107*(49), 21199-21204. |
| YieldMainEffect2009 | Yield was recorded as dry seed weight in grams. Main effect estimated from two walk-in growth chambers (AR-916, Percival Scientific) that were programmed to cycle the local climates every 5 min from the simulated weather files. The simulated climates were generated by using SolarCalc (35) with sunrise and sunset, light spectrum, temperature, and relative humidity programmed to cycle throughout the day and the season according to 1975?2000 averages (Fig. S5). One chamber was simulating Spain (latitude 41.72091, longitude 2.957075) starting from March 1, and the other chamber was simulating Sweden (latitude 55.71226, longitude 13.207352) starting from May 1st on the first day when plants were put into the chambers. | 2 | yield trait (TO:0000387) | 454 | Li, Y., Huang, Y., Bergelson, J., Nordborg, M., & Borevitz, J. O. (2010). Association mapping of local climate-sensitive quantitative trait loci in Arabidopsis thaliana. *Proceedings of the National Academy of Sciences*, *107*(49), 21199-21204. |
| FruitNumber | Total number of fruits per plant at the end of reproduction (fruit ripening). | 31 | yield trait (TO:0000387) | 409 | Vasseur, F., Exposito-Alonso, M., Ayala-Garay, O. J., Wang, G., Enquist, B. J., Vile, D., ... & Weigel, D. (2018). Adaptive diversification of growth allometry in the plant Arabidopsis thaliana. *Proceedings of the National Academy of Sciences*, *115*(13), 3416-3421. |
| GrowthRate | Average growth rate during the whole life cycle (final dry mass / lifespan, mg d-1). | 31 | shoot system growth and development trait (TO:0000928) | 420 | Vasseur, F., Exposito-Alonso, M., Ayala-Garay, O. J., Wang, G., Enquist, B. J., Vile, D., ... & Weigel, D. (2018). Adaptive diversification of growth allometry in the plant Arabidopsis thaliana. *Proceedings of the National Academy of Sciences*, *115*(13), 3416-3421. |
| root_length_day003 | Root length was scored 3 days post germination | 44 | shoot system growth and development trait (TO:0000928) | 231 | Satbhai, S. B., Setzer, C., Freynschlag, F., Slovak, R., Kerdaffrec, E., & Busch, W. (2017). Natural allelic variation of FRO2 modulates Arabidopsis root growth under iron deficiency. *Nature communications*, *8*(1), 15603. |
| Trichome_stem_length | Trichome stem length (abbreviated as stem length in the article) measured using the NIS elements software with manual structure identification; from photographs taken in the UV autofluorescence channel; average from at least 25 trichomes from 3 dried leaves of 3 plants. | 126 | plant [organ morphology trait](https://bioportal.bioontology.org/ontologies/PTO/?p=classes&conceptid=http%3A%2F%2Fpurl.obolibrary.org%2Fobo%2FTO_0000736) (TO:0000836) | 305 | Togninalli, M., Seren, Ü., Freudenthal, J. A., Monroe, J. G., Meng, D., Nordborg, M., ... & Grimm, D. G. (2020). AraPheno and the AraGWAS Catalog 2020: a major database update including RNA-Seq and knockout mutation data for Arabidopsis thaliana. *Nucleic acids research*, *48*(D1), D1063-D1068 |

### **Supplementary Table S2**: Prediction accuracy of each model across the 36 traits tested.

The following tables summaries the accuracy of each model for each trait analysed in the study

| **Trait** | **Ridge** | **Lasso** | **ElasticNet** | **RandomForest** | **MLP** |
| --- | --- | --- | --- | --- | --- |
| **herbavore_resistance_G2H3B** | **0.5057** | 0.5011 | 0.462 | 0.4775 | 0.4057 |
| **herbavore_resistance_G2H4P** | **0.4091** | 0.3727 | 0.3693 | 0.3744 | 0.173 |
| **herbavore_resistance_G2P** | 0.3767 | 0.3768 | 0.3817 | **0.4409** | 0.2581 |
| **herbavore_resistance_G3B** | **0.3153** | 0.2446 | 0.2736 | 0.3105 | 0.0816 |
| **herbavore_resistance_G3HP** | 0.3638 | **0.5089** | 0.4429 | 0.4963 | 0.4151 |
| **herbavore_resistance_G4MSB** | 0.4601 | 0.4488 | 0.4391 | **0.4772** | 0.4247 |
| **herbavore_resistance_G4P** | 0.2389 | **0.2294** | 0.2354 | 0.1895 | 0.0383 |
| **herbavore_resistance_G5MSP** | 0.2575 | 0.2706 | 0.2609 | **0.2892** | 0.182 |
| **study_16_Cd111** | 0.0489 | 0.2562 | 0.2379 | **0.282** | 0.0208 |
| **study_16_Mo98** | 0.1323 | 0.1072 | 0.2389 | **0.3385** | -0.0646 |
| **study_1_FLC** | **0.3386** | 0.1774 | 0.2308 | 0.2406 | -0.0967 |
| **study_1_FRI** | 0.0678 | 0.158 | 0.1802 | **0.2203** | 0.1795 |
| **study_4_M130T666** | 0.1195 | 0.5827 | 0.5609 | **0.6067** | 0.2543 |
| **study_12_FT10** | **0.6717** | 0.655 | 0.6532 | 0.626 | 0.6198 |
| **study_1_0W** | **0.5302** | 0.434 | 0.4298 | 0.4244 | 0.4503 |
| **study_1_8W_GH_FT** | **0.486** | 0.4093 | 0.3943 | 0.3912 | 0.319 |
| **study_1_FT22** | **0.3933** | 0.2731 | 0.286 | 0.1908 | 0.3089 |
| **study_1_FT_Field** | **0.5274** | 0.4618 | 0.4635 | 0.4111 | 0.0498 |
| **study_1_SDV** | **0.4115** | 0.2857 | 0.3145 | 0.2824 | 0.08 |
| **study_2_DTFmainEffect2009** | **0.5908** | 0.6225 | 0.5783 | 0.5082 | 0.5301 |
| **study_126_Trichome_stem_length** | **0.2694** | 0.1977 | 0.201 | 0.2675 | -0.0011 |
| **study_1_Width_10** | 0.0948 | **0.2178** | 0.0979 | 0.1376 | 0.0434 |
| **study_27_MeanTRL_C** | **0.2001** | 0.0918 | 0.1425 | 0.1562 | 0.0679 |
| **study_30_MeanTRL_CK** | **0.2597** | 0.2436 | 0.1896 | 0.15 | 0.0196 |
| **study_38_CL** | **0.4679** | 0.3708 | 0.4025 | 0.4168 | 0.2484 |
| **study_38_RL** | **0.5364** | 0.5017 | 0.5159 | 0.4697 | 0.4193 |
| **study_44_root_length_day003** | **0.2093** | 0.0273 | 0.1322 | 0.1612 | -0.0527 |
| **study_19_GR21** | **0.5592** | 0.5307 | 0.5057 | 0.5506 | 0.5446 |
| **study_1_Storage_28_days** | **0.3515** | 0.4454 | 0.2952 | 0.3384 | 0.3312 |
| **study_1_Storage_56_days** | **0.291** | 0.2674 | 0.2152 | 0.2309 | 0.1564 |
| **study_1_8W_GH_LN** | **0.4699** | 0.3025 | 0.2796 | 0.2386 | -0.1364 |
| **study_1_LN10** | **0.4852** | 0.4823 | 0.396 | 0.3809 | 0.0918 |
| **study_1_LN16** | **0.5272** | 0.4546 | 0.4444 | 0.388 | -0.0795 |
| **study_31_GrowthRate** | **0.3765** | 0.3289 | 0.3232 | 0.3075 | 0.169 |
| **study_2_YieldMainEffect2009** | **0.2256** | 0.156 | 0.1932 | 0.1996 | 0.0977 |
| **study_31_FruitNumber** | **0.2946** | 0.2675 | 0.2512 | 0.2699 | 0.0516 |

### **Supplementary Figure S1**: Manhattan plots of the GWAS for the 36 traits analysed.

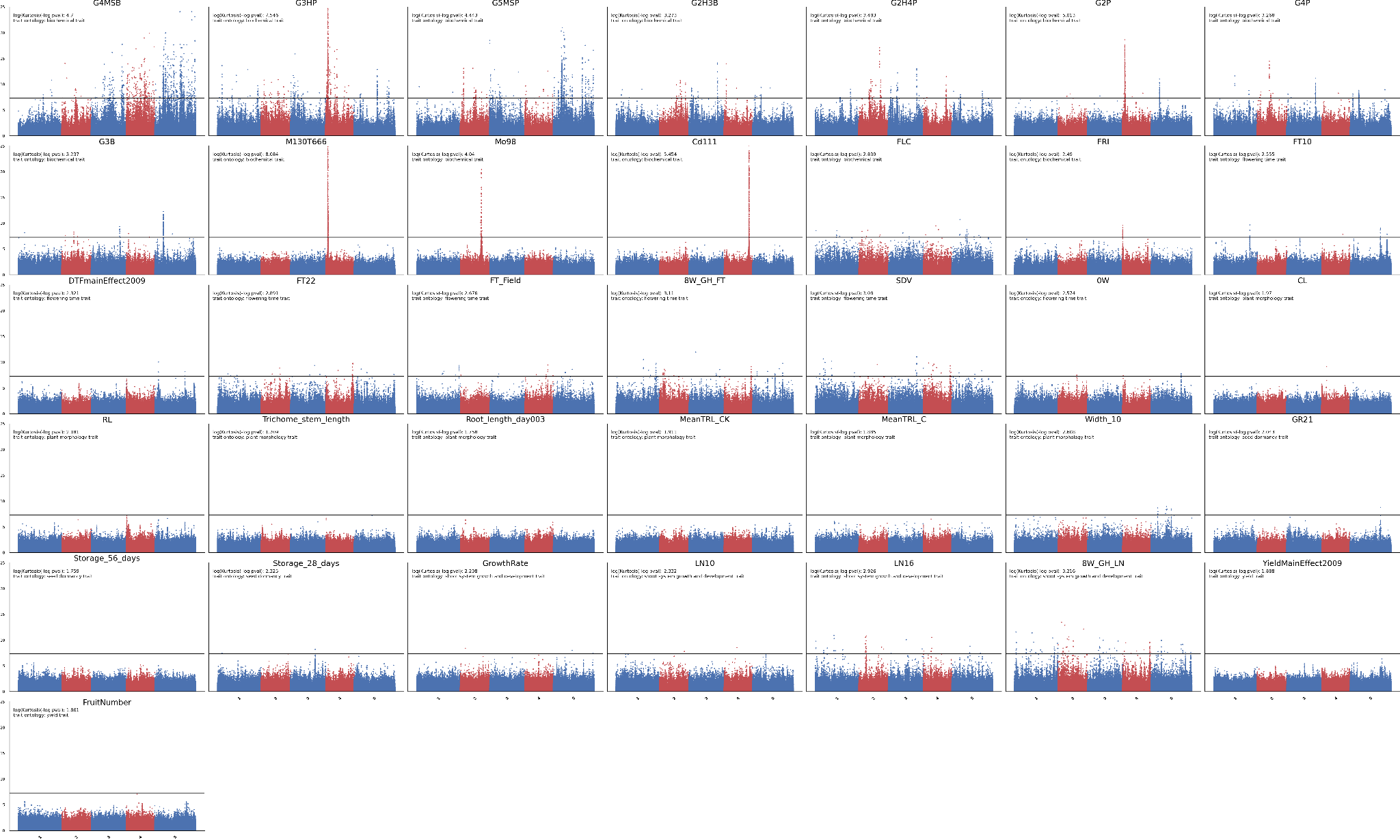
